# Intestinal crypts recover rapidly from focal damage with coordinated motion of stem cells that is impaired by aging

**DOI:** 10.1101/218685

**Authors:** Jiahn Choi, Nikolai Rakhilin, Poornima Gadamsetty, Daniel J. Joe, Tahmineh Tabrizian, Steven M. Lipkin, Derek M. Huffman, Xiling Shen, Nozomi Nishimura

**Affiliations:** Meinig School of Biomedical Engineering, Cornell University, Ithaca, New York, 14853, USA.; Biomedical Engineering, Duke University, Durham, North Carolina, 27708, USA.; Gastroenterology, Weill Cornell College of Medicine, New York, New York, 10065, USA.; Molecular Pharmacology & Medicine, Albert Einstein College of Medicine, Bronx, New York, 10461, USA.

## Abstract

Despite the continuous renewal and turnover of the small intestinal epithelium, the intestinal stem cell niche maintains a ‘soccer ball-like’, alternating pattern of stem and Paneth cells in the crypt. To study the robustness of the niche pattern, we used intravital two-photon microscopy in mice with fluorescently-labeled Lgr5+ intestinal stem cells and precisely perturbed the mosaic pattern with femtosecond laser ablation. Ablation of one to three cells initiated rapid motion of niche cells that restored the alternation in the crypt pattern within about two hours without any cell proliferation. Crypt cells then performed a coordinated dilation of the crypt lumen, which resulted in peristalsis-like motion that forced damaged cells out of the niche. Crypt cell motion was reduced with inhibition of the ROCK pathway and attenuated with old age, and both resulted in incomplete pattern recovery. This suggests that in addition to proliferation and self-renewal, motility of stem cells is critical for maintaining homeostasis. Reduction of this novel behavior of stem cells could contribute to disease and age-related changes.

## INTRODUCTION

The rapid regeneration of the intestinal epithelium is enabled by fast-cycling Lgr5+ intestinal stem cells (ISCs) crowded into the base of the crypt^1-3^. ISCs are not only limited in number and location, but also arranged in a specific pattern. ISCs expressing Lgr5 are intercalated between Paneth cells, which are secretory cells with antibacterial functions. This organization results in a ‘soccer ball-like’, mosaic pattern in which Lgr5+ ISCs form a continuous network that surrounds each Paneth cell^4^. In the healthy crypt, this alternating pattern is persistent despite frequent cell division and migration^2,5^, but the dynamics of how this architecture is maintained is unknown.

To investigate the robustness of the patterning and its maintenance *in vivo*, we ablated individual cells in the crypt with high-pulse-energy femtosecond (fs) laser ablation, and imaged the real-time dynamics of recovery with multiphoton microscopy^6,7^. Such accurate manipulation is not achieved by current methods of radiation or chemical treatment, or genetic ablation of specified lineages^8,9^. Surprisingly, after ablation of a small number of cells, migration of neighboring cells was sufficient to reestablish cellular contacts and the alternating pattern in the crypt base within hours, before any cells divided. In addition, we observed coordinated motion of the cells at the edge of the crypt base that expelled debris out towards the lumen. The repair movements were impaired by both ROCK inhibition and aging, highlighting the importance of this dynamic response for the integrity of the niche.

## RESULTS

### Two-photon microscopy enables visualization of alternating pattern in the crypt

To understand how the niche spatiotemporally maintains homeostasis *in vivo*, we modified previously demonstrated abdominal imaging windows^10,11^ with a 3D printed insert that sits under a portion of the small intestine to minimize movement without blocking normal digestive functions (Fig. 1a, b). For imaging, animals were anesthetized and the window frame was fixed with a clamp under the objective of a two-photon microscope with ablation capabilities (Fig. 1c). Lgr5+ crypt cells were visualized in mice expressing green fluorescent protein driven by the Lgr5 promoter (Lgr5-GFP)^3^, while Paneth cells appeared dark (Fig. 1d). The vasculature was labeled by retro-orbital injection of Texas Red dextran (Fig. 1d). Blood vessels were stable over weeks, enabling us to use them as a roadmap to image the same areas repeatedly (Suppl. Fig. 1a). We injected Hoechst under the window to label nuclei. Time-lapse images show that the pattern of GFP and the number of Hoechst-labeled nuclei within a crypt was stable over the course of several hours (Fig. 1e, Suppl. Fig. 1b).

**Figure 1.**
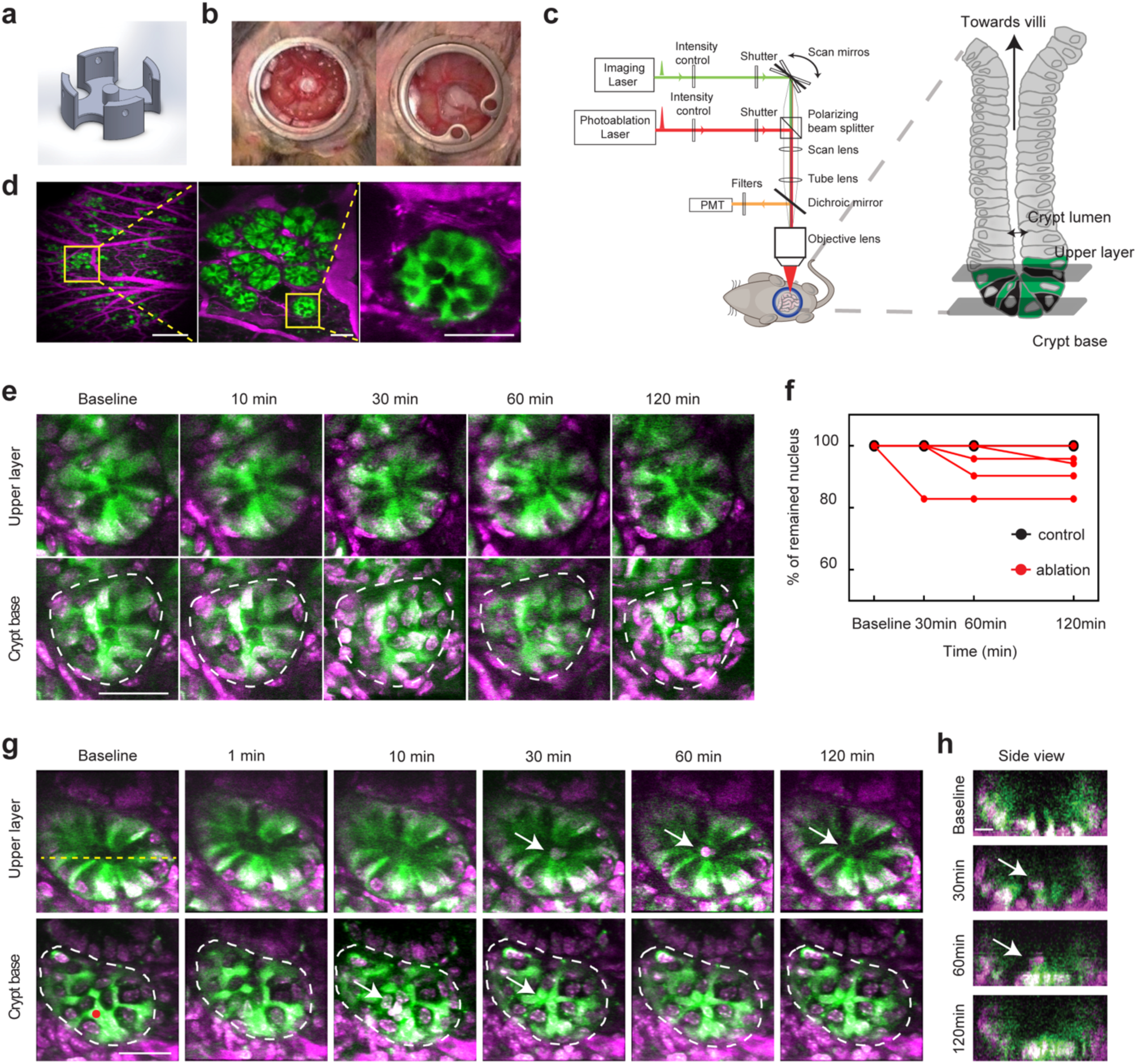
Rapid clearance of local damage after femtosecond laser ablation in a mouse small intestinal crypt. (**a**) Schematic of 3D-printed insert that supports intestine in abdominal window. (**b**) Photos of abdominal window before (left) and after (right) closing with cover glass in a mouse. (**c**) Schematic (modified from Chen et al. #19) of two-photon microscope showing two different laser paths used for both imaging and ablation. (**d**) In vivo two-photon microscopy images of a crypt at different magnifications in Lgr5-GFP mice expressing GFP in stem cells at the crypt base (green). Vessels are labeled with injected Texas Red dextran (magenta). Yellow boxes indicate magnified areas. Scale bars: 500 µm (left), 50 µm (middle and right). (**e**) Time-lapse images showing two different imaging planes in a crypt over 2 hours. Green indicates GFP. To label nuclei, Hoechst (magenta) was injected topically. Dashed white lines indicates the border of the crypt base. Scale bar: 30 µm. (**f**) Number of nuclei in crypt base after ablation (red) and control (black). (**g**) Time-lapse images of femtosecond laser ablation of one Lgr5-GFP cell in a crypt at two image planes. Red dot indicates position of ablation laser focus. White arrow indicates cellular debris from the ablation which moved from crypt base towards the villi. Scale bar: 30 µm. (**h**) Side view at line indicated in (**g**). Scale bar: 10 µm.

### Cells damaged by femtosecond laser ablation are expelled from the crypt base

Cells were ablated selectively during imaging with photodisruption^12,13^ by pulses from a Ti:Sapphire regenerative amplifier. The damage was largely confined to the focal volume while neighboring cells and adjacent crypts were not affected (Suppl. Fig. 1c,1d). In contrast, attempted ablation with the imaging beam at high power resulted in damage in a large region (Suppl. Fig. 1e). We first targeted a single Lgr5+ ISC in the crypt base.

The GFP fluorescence from the targeted cell quickly dissipated, but nuclear labeling was still detected at the ablated site. Over the next 10-30 minutes, the nucleus of the ablated ISC disappeared from the base of the crypt and moved through the crypt lumen in the direction of the villi. Nuclei of the remaining cells appeared intact for the duration of the imaging time, up to 2 hours after ablation (Fig. 1g, h; Suppl. Fig. 1f). The ablation debris, still labeled with Hoechst, then gradually passed through the lumen until it was beyond the 50-µm field of view. Ablation debris always moved up towards the villi and never towards the lamina propria of the intestine (74 crypts). Once the damaged cells were pushed out into the lumen, the number of remaining Hoechst-labeled nuclei at the base of the crypt did not change. In adjacent control crypts without ablation, the number did not change during 2 hours (Fig. 1f).

### Alternating pattern of intestinal stem and Paneth cells is rapidly restored by cell motion

Laser ablation of a Paneth cell resulted in a disruption and recovery similar to ablation of Lgr5+ ISCs (Fig. 2a, Suppl. Fig. 2b). At the base of the crypt, neighboring Lgr5+ ISCs moved towards the damaged area and restored a spatial configuration in which Paneth cells were separated from other Paneth cells by Lgr5-GFP cells within 2 hours. At 24 hours, the crypt was intact and had an alternating cell pattern (Fig. 2a, control crypt in Suppl. Fig. 2a). More extensive disruption was produced by ablating three ISCs (Fig. 2b) or three Paneth cells (Suppl. Fig. 2c), but these crypts were also able to re-establish the intercalating pattern by rearrangement within 2 hours. We quantified the change in the alternating ISC and Paneth cell pattern between baseline and 1 day after ablation of 1 to 3 ISCs or Paneth cells by calculating a rearrangement score that describes the change in alternating positions between ISCs and Paneth cells (Fig. 2c). Ablated crypts showed greater disparity from the baseline pattern than without ablation (Fig. 2d).

**Figure 2.**
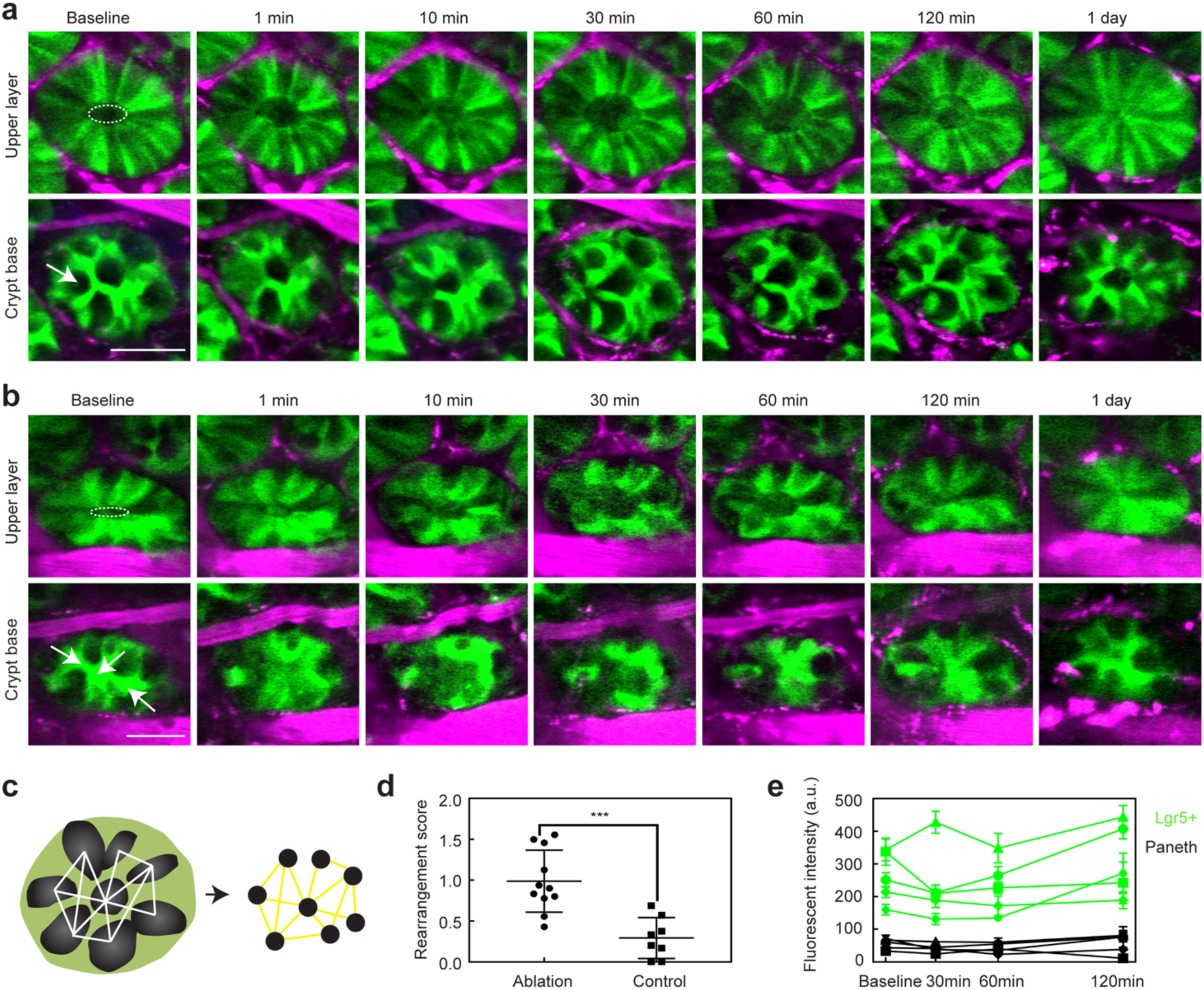
Perturbation of the crypt base pattern was followed by coordinated motion which restored the crypt base pattern and removed the debris from ablation. (**a**) Time-lapse images of ablation of a single Paneth cell showing two imaging planes. White arrow indicates the position of the ablation laser focus. In the upper layer, the white dashed ellipse indicates the circumference of crypt lumen at baseline. Scale bar: 30 µm. (**b**) Time-lapse images of ablation of multiple stem cells by focusing laser in three positions indicated by white arrows. White dotted ellipse shows the inner circumference at the baseline. Scale bar: 30 µm. (**c**) Schematic showing the strategy for describing the alternating pattern using network graphs. Each Paneth cell is a node (black). Edges were defined when nearest Paneth cells were separated by Lgr5+ ISCs. (**d**) Quantification of change in alternating ISC and Paneth cell pattern at crypt base using rearrangement score based on adjacency matrices calculated from networks defined in (**c**). *** p=0.0003, unpaired t-test. (**e**) Mean and standard deviation of fluorescent intensity from regions manually chosen in Lgr5+ ISCs and Paneth cells over time.

We next aimed to determine whether this observed pattern change resulted from loss of Lgr5+ ISC identity as reported by Lgr5-GFP, so we measured the intensity of fluorescence from GFP near the ablated site as a function of time after ablation of 1-3 cells (Fig. 2e). Although there were small fluctuations, the intensity of bright, Lgr5+ ISCs and dark, Paneth cells remained distinct. As a positive control for loss of GFP signal of Lgr5+ ISCs, we applied dibenzazepine (DBZ), which inhibits Notch signaling and ISC self-renewal, under the window 2 hours before imaging, then ablated cells at a crypt. DBZ treatment decreased baseline intensity of Lgr5-GFP fluorescence, and the Lgr5+ ISCs and Paneth cells were no longer distinguishable by intensity after ablation (Suppl. Fig. 2d).

### Dilation and contraction of the crypt lumen pushes debris towards villi

During the rearrangement of the cells located at the crypt base after ablation, the diameter of the crypt lumen, 15-20 µm above the crypt base, enlarged over 2 hours (Fig. 2a, b). However, the outer diameter of the crypt stayed relatively constant during this time. To label damaged cells, we injected propidium iodide (PI, retro-orbital) immediately before imaging, which enters cells only if the cellular membrane is disrupted, and emits red fluorescence when bounds to DNA^14^. In unperturbed crypts, PI labeling was rarely observed, but red fluorescence was readily detected within several minutes at the targeted site following ablation. PI labeling clearly demonstrated that damaged cells were expelled towards the crypt lumen within an hour and then upward through the lumen toward the villi (Fig. 3a). The motion of the PI-labeled debris coincided with the crypt lumen dilation.

**Figure 3.**
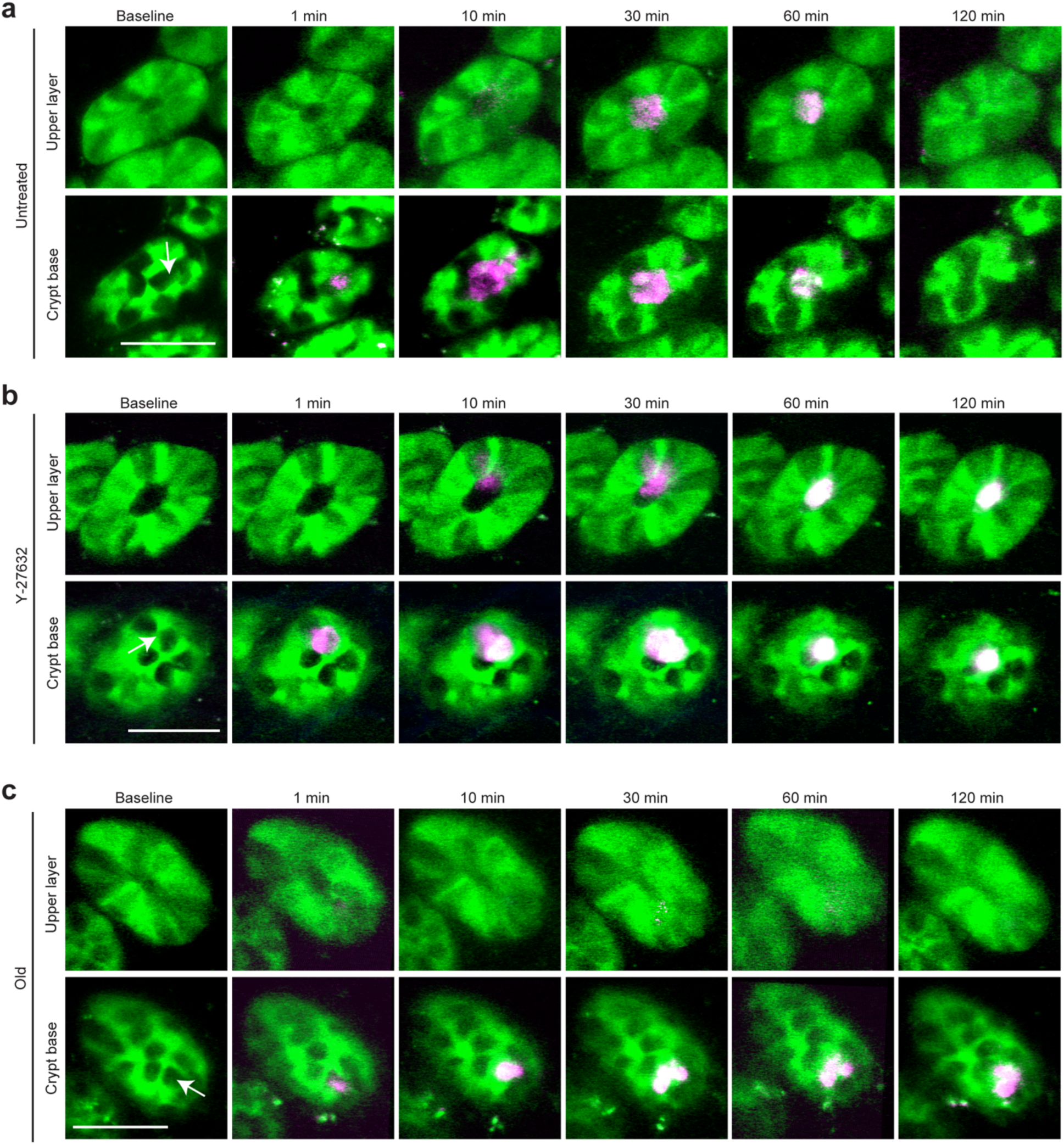
Cell motions after ablation are reduced after inhibition of ROCK signaling and in aged mice. (**a**) Time-lapse images after ablation of one Paneth cell in crypt with Lgr5+ ISC (green) and propidium iodide (PI) (magenta) indicating damaged cells in a young mouse (6-month old). White arrow indicates the position of ablation laser focus. Scale bar: 50 µm. (**b**) Time-lapse images of ablation of one Lgr5+ ISC in crypt in an inhibitor-treated, young mouse (4-month old). To inhibit the cellular motility, Y-27632, was topically administered. A white arrow indicates the focus of ablation laser. Scale bar: 50 µm. (**c**) Time-lapse images of ablation of one Paneth cell in an aged mouse (17-month old) with PI injection. A white arrow indicates the focus of laser ablation. Scale bar: 50 µm.

### Crypt cell motion is dependent on the ROCK pathway

To test whether this process was due to actin-mediated cell motion, as opposed to a passive reaction to external forces, we inhibited the ROCK pathway with topical application of Y-27632^15^. ROCK inhibition did not noticeably change the baseline patterns, but reduced motion of the crypt cells, which resulted in incomplete crypt base pattern at 1 day after ablation (Suppl. Fig. 3). The PI-labeled damaged cells remained at the base and elimination of the debris was incomplete after two hours (Fig. 3b). ROCK inhibition also affected crypt lumen dilation. In the crypt treated with Y-27632, the lumen did not change in size and the debris was detected in the lumen for an extended time period.

### Cellular motion is impaired in aging crypts

Functional decline is a hallmark of aging in multiple tissues, including the gut, and this process is thought to be driven in part by deterioration in resident stem cell function^16,17^. Therefore, we tested how the aged niche cells respond to local damage by comparing young (2-6 months) with old (17-23 months) Lgr5-GFP mice. After ablation of a single Paneth cell, the aged crypt showed a response that was similar to an inhibitor-treated crypt. Cells at the crypt base did not show noticeable rearrangement after ablation, and PI-labeled damaged cells were remained at the base of the crypt for 2 hours, while at the upper layer, the crypt lumen had minimal dilation (Fig. 3c).

### Area of damage is increased in aged and ROCK-inhibited crypts

To quantitatively compare damage in young animals with and without Y-27632, and aged animals, we used the area of PI labeling at and just above the base after ablation of a single cell (both ablation of ISCs and Paneth cells were included). Immediately after ablation, the PI-labeled area at the base was the same across all groups, but the old group increased the most over time. In young, untreated mice, the PI-labeled area at and just above the crypt base increased and then decreased as the debris moved towards the villi (Fig. 4a, b). The peak value of the area was slightly delayed at the upper plane relative to the crypt base. Neither the young, treated group nor the old group showed decrease in PI area over two hours. In the old mice, the PI-labeling at the base was consistently larger than both young groups. In the upper plane, the PI-area in the old group was similar to the young, treated group. A linear regression to the PI area in the base of the crypt between 30 and 240 minutes (time after peak area) showed that only the young, untreated group had a negative slope (R^2^ = 0.99, p = 0.0052), while in young, treated and old mice the slopes suggest PI area remains constant. At the upper level, the young, untreated group was also the only group to have a negative slope, but this did not reach significance.

**Figure 4.**
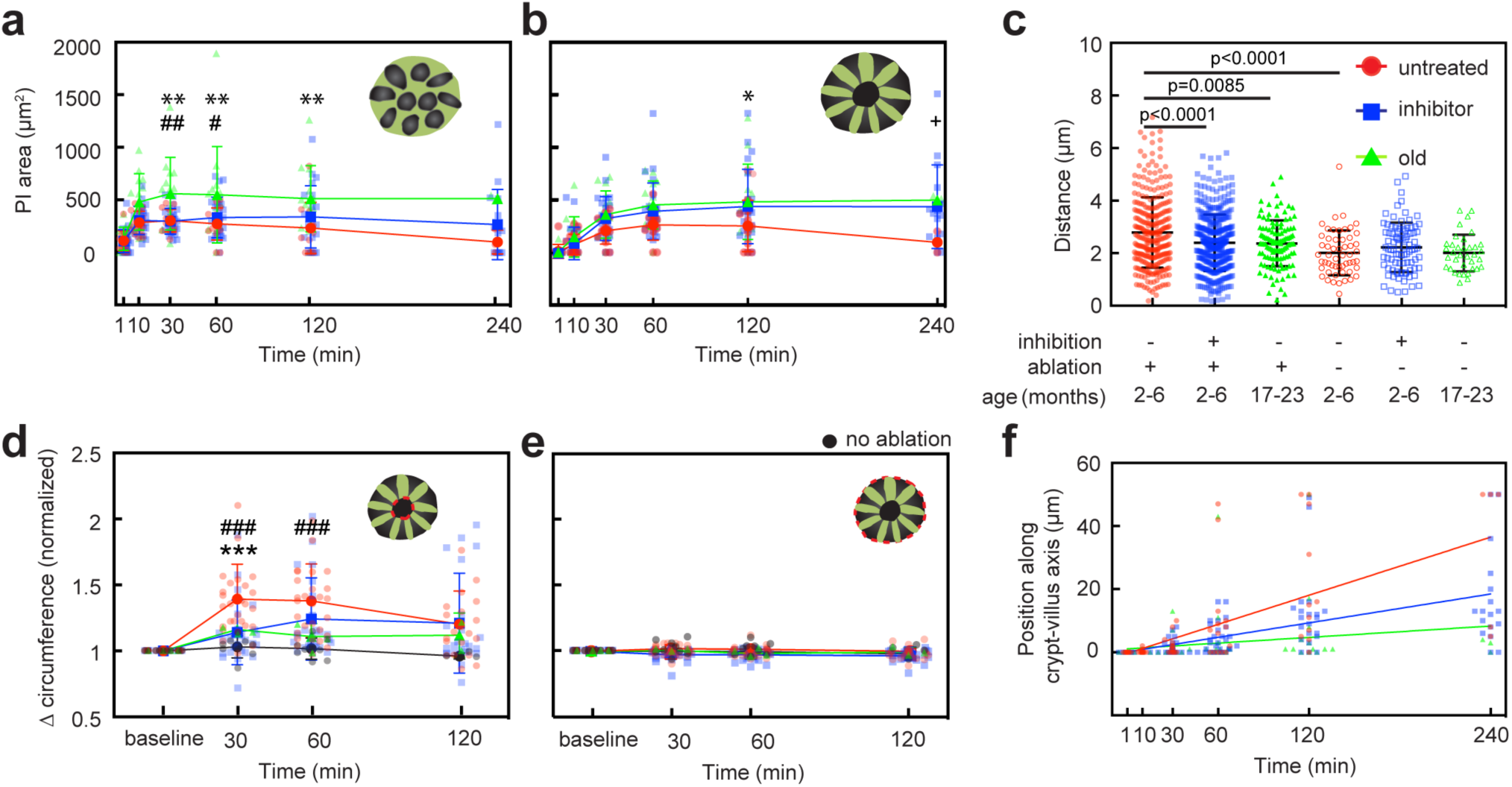
Comparisons of damage and crypt cell motion after single cell ablation in young mice with and without Y-27632, and aged mice. (**a-f**) Comparison of crypt cell behavior in untreated, inhibited, and old mice. Red color indicates untreated, young mice, blue color is young, Y-27632 treated mice, and green color indicates old mice with no treatment. At each measurement time, dots are spaced for visual clarity. (**a-b**) Area labeled with propidium iodide (PI) after ablation at the crypt base (**a**) and at the upper layer (**b**). Mean and SD are plotted at each time point and dots represent individual crypts. * indicates comparison between untreated, young mice and old mice, # between Y-27632 treated, young mice and old mice, and + between untreated young mice and Y-27632 treated, young mice. **, ##, ++ indicates p<0.01, and *, #, + indicates p<0.05 (2-way ANOVA with Tukey’s multiple comparisons). (**c**) Sum of distance moved by Paneth cells at the crypt base between baseline and one hour after ablation. Mean and SD are plotted with dots representing individual Paneth cells. Significance calculated with ANOVA with Tukey’s multiple comparisons. (**d-e**) Measurement of inner (**d**) and outer circumference (**e**) at the crypt lumen at each time point after ablation. Each value was normalized to the baseline. Mean and SD are plotted with dots representing individual crypts. Black indicates measurements in control crypts adjacent to ablated crypts. * indicates comparison between untreated, young mice and Y-27632 treated, young mice, and # indicates comparison between untreated, young, ablated mice and untreated, young, unablated mice. ***, ### indicates p=0.0003 (2-way ANOVA with Tukey’s multiple comparisons). (**f**) Position of cellular debris along the crypt lumen. Dots represent individual crypts at each time point. Lines represent linear regression.

### Both rearrangement and peristalsis are reduced by age and ROCK inhibition

To compare the amount of rearrangement of base cells after ablation quantitatively, we measured the distance of Paneth cell movement summed over time points for the first one hour after ablation (Fig. 4c, Suppl. Fig. 2e). In young, untreated mice, Paneth cells in the damaged crypts moved on average about 135% further than those in adjacent, unablated, control crypts (Fig. 4c). After ablation, Paneth cells in young, Y-27632-treated mice and old mice moved only about 86% as far as in young, untreated mice. ROCK inhibition and aging decreased dilation of the crypt lumen relative to the young, untreated group (Fig. 4d). After ablation, the inner circumference of untreated crypts in young mice increased up to 142% of baseline at one hour and then decreased back towards baseline by 2 hours. In adjacent, unablated crypts, there was no change in either the inner or outer diameter. Y-27632 treatment reduced the increase of the inner circumference to only 125% of baseline and in old animals, the inner circumference only increased up to 110% of baseline. The outer circumference remained unchanged in all groups (Fig. 4e).

We next measured the average speed of debris clearance by tracking the position of PI-labeled cellular debris as it moved toward the villi after ablation (Fig. 4f). Untreated crypts in young animals showed the fastest speed, estimated from the slope of the position measured over 4 hours at 14 µm/hour, while inhibitor-treated and aged crypts had markedly slower elimination speed (7 µm/hour and 1 µm/hour, respectively). Using linear regression analysis, the slope of the untreated group showed a significant difference from Y-27632 treated, and the old group (R^2^=0.9665, p=0.0023 for untreated vs inhibitor, p<0.0001 for untreated vs old). Collectively, these data suggest that the recovery process accompanied by coordinated cellular motion is a regulated process whose dynamics are impaired with aging (Fig. 5).

**Figure 5.**
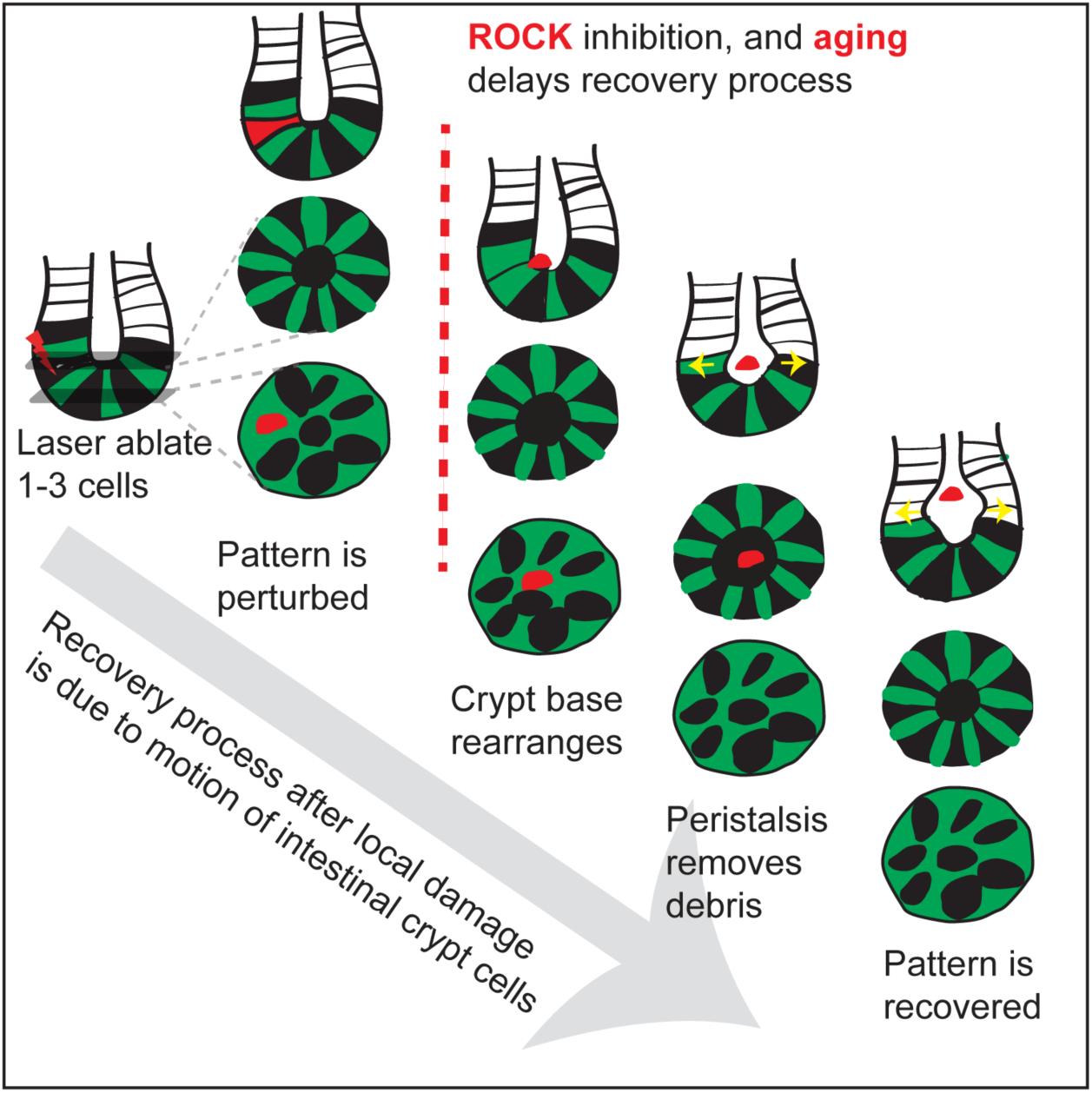
Coordinated cellular motion facilitates rapid recovery of crypt patterns after focal damage. Schematic summarizing the recovery process after pattern disruption by ablation of crypt cells with laser ablation. Orchestrated motion of crypt cells restores the alternating pattern while pushing the cellular debris out toward the villi. These motions are attenuated with ROCK inhibition and aging resulting in a disrupted crypt pattern.

## DISCUSSION

How and why the crypt niche maintains the precise spatial arrangement of Paneth and Lgr5+ cells during the constant division of ISCs is still a perplexing question^18-21^. In this study, *in vivo* time-lapse imaging and femtosecond laser photodisruption revealed that the response to localized ablation of a few crypt cells is immediate. Surrounding cells quickly mobilize to extrude the debris. Cell migration fills the vacancy in the epithelium and restores the alternating pattern within hours, well before stem cells proliferate. The debris from ablated cells is pushed out of the plane of the crypt base, in a manner similar to the removal of damaged cells in cultured epithelial monolayers, chick embryonic epithelium, *Drosophila* and developing brain of *Xenopus laevis*^22-25^. The speed of this expulsion was reduced by a ROCK inhibitor, suggesting that the mechanism in crypts could be similar to these other systems, in which dying cells activated neighboring cells to form a contractile actin ring. Even when different numbers of the two cells types were ablated (Fig. 2a, b; Suppl. Fig. 2b,c), the remaining cells still migrated to restore alternation in the cell pattern within 2 hours. The final patterns differed from those prior to ablation, suggesting that the arrangement is defined by interactions between the Paneth and stem cells, rather than dictated by the underlying mesenchymal cells. Signaling, such as Wnt/Notch, involved in maintaining stemness may also play a role in regulating this dynamic positioning^4,19,26,27^. Lgr5-GFP labeling remained constantly bright (Fig. 2e), suggesting that the process of rearrangement does not require loss or gain of the stem cell phenotype. Although conversion of other cells to the stem cell phenotypes is known to occur with certain types of injury and stimuli^8,28^, it is likely that the death of only a few cells with laser ablation is insufficient to trigger such a response.

In the intestinal crypt, the stem cells are at the end of a narrow tube, so that even after the expulsion from the crypt base, the cellular debris is trapped near the stem cells. The crypt cells performed a second type of motion, a peristalsis-like dilation of crypt lumen that appears to push the debris up the crypt towards the intestine interior. The dilation of the crypt lumen is likely crucial for effective extrusion of debris from the crypt base because it coincided with the arrival of debris in the crypt lumen. The peak dilation is earliest at the base of the crypt and later towards the villi, consistent with peristaltic pumping (Fig. 4a, b). The outer diameter does not change (Fig. 4e), suggesting that the changes of inner circumference resulted from contraction of crypt cells rather than from forces exerted by the myofibroblasts surrounding the crypt or muscle cells of the muscular mucosae. Such a coordinated response has been observed in neural progenitor cells in the developing neuroepithelium after laser ablation^23^, but has not been previously observed in adult stem cells. Activation of ROCK is a fundamental to regulate actin cytoskeletal activities and, although inhibition of ROCK pathway is a promising therapeutic target to alleviate with fibrosis in various organs^29,30^, consistent inhibition of such pathway could impair integrity of epithelium in pathophysiological conditions^31,32^.

Sporadic cell death, similar in sparsity to the carefully-controlled damage from femtosecond laser ablation, occurs in several biological processes. Most of the apoptosis in the intestinal epithelium that maintains constant cell numbers occurs at the villi, but occasional apoptosis does occur in the crypt^33,34^, suggesting that the rearrangement and peristalsis may play a role in normal function. In infection or IFN-γ stimulation, Paneth cells degranulate and then disappear from the stem cell niche^35-37^. In such instances, it is not only critical that fast cycling Lgr5+ ISCs can renew the epithelium with functioning cells, but that dead and damaged cells can be quickly and efficiently cleared from the cryptal microenvironment. Impaired clearance of damage in the stem cell niche can result in unnecessary inflammation^38^. Therefore, rapid removal of dying cells is important not only for pattern recovery, but also for limiting exposure to a dangerous environment.

Time-lapse imaging showed that the alternating pattern was restored within several hours, suggesting that the niche starts the repair process without waiting for ISCs to divide to replace the lost cells. Nevertheless, the stem cells will eventually divide to restore the total number of cells. It has been previously shown that the “central” ISCs have a survival advantage over border ISCs, which may be pushed into the trans-amplifying compartment to start differentiation^39^. Future studies using intravital abdominal imaging may reveal how cell proliferation and plasticity contributes to the repopulation of the crypt. These novel observations could shed insights into previously unappreciated contributors to intestinal function and dysfunction in the context of intestinal diseases, infection and normal aging.

Aging is known to alter the number, rejuvenative capacity and senescence of stem cells^16,40^, but we have identified an additional function of ISC and Paneth cell that declines with aging, their motion dynamics in response to a perturbation. In other types of cells, aging is associated with a decrease in cellular function that results in less effective cytoskeletal remodeling and slower motion^41^. Extrinsic factors, such as increased tissue stiffness with aging, could in part underlie this alteration in cell contractility and motion^42-44^. Here, we found that the motion of neighboring cells was dramatically impaired, and cell debris ‘lingered’ in the crypts of old animals. Importantly, the observation that the aged crypt harbors reduced capacity to expel dead cells, thereby aggravating the local inflammatory response, could have important implications for the maintenance of intestinal homeostasis.

## MATERIALS AND METHODS

### Animals

To visualize stem cells in the crypt we used *Lgr5* ^*tm1(cre/ERT2)Cle*^/*J* mice (Lgr5-GFP) bred at Cornell University from breeders originally purchased from the Jackson Laboratory (Stock No. 008875). Some Lgr5-GFP mice were crossed with *Gt(ROSA)26Sor^tm1(EYFP)Cos/J^* mouse purchased from the Jackson Laboratory (Srinivas, 2001 #108; Stock No. 006148). Other Lgr5-mice were bred with C57BL/6J mice (Stock No. 000664). No clear differences in labeling or phenotype were observed between these two breeding schemes so all data has been pooled. For aging studies, old and young mice were bred at Albert Einstein College of Medicine, and shipped to Cornell where they were housed for at least 3 weeks before imaging studies. No differences were apparent between the young mice from the Cornell and Einstein colonies, so data from both colonies were combined. All animals were housed at standard temperature (72 °F) and humidity-controlled conditions under a light cycle of 14 light and 10 dark hours with most imaging performed during the light hours. Mice were provided *ad libitum* access to water and a regular chow diet (Teklad 7912). Age and sex of subjects in each graph is provided in the supplementary table. All animal care and experimental procedures were approved by the Institutional Animal Care and Use Committee of Cornell University and were conducted in accordance with relevant guidelines and regulations.

### Abdominal window preparation

Mice were anesthetized with 4% isoflurane in oxygen, and maintained at ~2%. At the beginning of the surgery, animals received an anticholinergic, 0.05 mg per 100 g of mouse weight of glycopyrrolate or 0.005 mg/100 g of atropine subcutaneously, to assist in keeping the airways clear of fluid build-up. Body temperature was kept at 37 C using a heating blanket controlled by a rectal thermometer. All areas to be incised were shaved and cleaned with 70% ethanol, and betadine, and were numbed with a 100 µL subcutaneous injection of 0.125% bupivacaine. Eyes were covered with veterinary eye ointment to prevent drying. The animals were hydrated with subcutaneous injections of 5% glucose in saline for isotonic fluid replacement. Abdominal skin was removed in a circular shape to fit a titanium window frame (Brain Titanium Chamber Kit, 12 mm in inner diameter, APJ trading, CA). The frame was put on top of the abdominal muscle. The outer edge of the frame was covered by skin and secured with cyanoacrylate adhesive (Loctite 406, Henkel). The abdominal muscles within the inner circle were incised with scissors to expose the small intestine. Custom-designed scaffold to stabilize the intestine were 3D-printed (UPrint) out of ABSplus (SolidWorks file in Supplementary Materials). The insert was placed under the small intestine so that a portion of the small intestine looped around the center post, entering and exiting the insert at the same gap between the supports. The insert was sutured to the skin outside of the frame by threading sutures through the holes in the four supports. The window was closed with a 12-mm diameter cover glass held on by the frame’s retaining ring. Window and insert was placed in the abdominal area 1-1.5 cm below the rib cage. Dexamethasone (0.025 mg/100 g) and ketoprofen (0.5 mg/100 g) were injected intraperitoneally to help recovery after surgery and then daily for 2 days. Animals were returned to their home cage after recovery. Imaging was performed at least 24-48 hours after implantation to allow the position of the intestine in the window to settle and to minimize the effects of the surgery.

### In vivo two-photon excited fluorescence microscopy

Animals with abdominal window were imaged using a custom-built two-photon excited fluorescence (2PEF) microscope. All animals were anesthetized with isoflurane (1-2% in medical air) which was adjusted to maintain constant breathing rate during imaging time. To visualize vasculature, 50 µl of 2.5% Texas Red dextran (Molecular weight: 70,000; Thermo Fisher Scientific, NY) in saline was injected retro-orbitally. Images were acquired using a Ti:Sapphire laser (Chameleon; Coherent, Santa Clara, CA), with wavelength centered at 880 nm. Either a 20x, 1.0-NA (Ziess, Thornwood, NY) or 25x, 1.05-NA (Olympus) water-immersion microscope objective was used for imaging and ablation. A 4X, 0.28-NA objective lens (Zeiss, Thornwood, NY) was used for low-resolution mapping. A 494-nm bandpass filter with 41-nm bandwidth and a 641-nm bandpass filter with a 75-nm bandwidth was used for green fluorescent protein (GFP) and Texas Red dextran dye respectively. For Hoechst-labeled crypt imaging, we tuned the Ti:Sapphire laser to 830 nm. A 458-nm bandpass filter was used to detect Hoechst. Image stacks were acquired before and after laser ablation at 1.44 frames/s, with 1-µm spacing in the axial direction.

### Disruption of cellular contact using femtosecond laser photodisruption

Selective disruption of cells was performed using a low-repetition-rate, high-pulse-energy Ti:Sapphire regenerative amplifier with 50-fs pulse duration, 1-kHz repetition rate, and 800-nm central wavelength (Legend-USP; Coherent, Santa Clara, CA). A polarizing beamsplitter cube was used to introduce this beam into the 2PEF microscope so that the pulses were focused at the center of the imaging field and in the 2PEF imaging plane, enabling real-time monitoring (Fig. 1c). Laser energy incident on the cells was controlled by neutral density (ND) filters, and a fast mechanical shutter limited the number of pulses (2-5) incident on each cell. We started ablation with lower power by using highest ND filter, and then gradually increased the power until we saw the formation of a small hole in the cytoplasm (Suppl Fig. 1c). Incident laser energy to remove a single cell was typically about 50 nJ per pulse and did not exceeded 100 nJ.

### Addition of chemical compounds

To label all nuclei in the crypt base, Hoechst-33342 (Thermo Fisher, Watham, MA; 2mg/kg) was injected under the window 1 hour before imaging. To inhibit ROCK protein signaling, Y-27632 (Sigma Aldrich, St. Luis, MO; 5mg/kg mouse weight in saline) was injected under the window before the start of imaging. This dosage was selected as the highest dose which did not alter crypt structure or change GFP expression patterns over one day (data not shown). Saline (0.9 %) (Phoenix Pharmaceutics, Burlingame, CA) was injected for the sham-treated group. To label damaged cells, propidium iodide (Cayman Chemical Company, Ann Arbor, MI; 100 µl of 1 mg/ml in PBS) was injected retro-orbitally. Dibenzazipene (DBZ, Syncom, Netherlands), a small molecule gamma-secretase inhibitor, was dissolved in DMSO at final concentration of 30 µM, and locally injected into the submucosal layer of the small intestine through the implanted abdominal window on mice (15 µM/kg) 2 hours before imaging.

### Image processing

All images were processed using ImageJ. Frames that contained abrupt movement from respiration or peristalsis were deleted manually (approximately 20% of frames). To enhance signal/noise ratio, we applied a median filter with 1~2 pixel radius depending on the noise. Images within the stack was registered using a macro function called Stackreg^45^. For quantitative analysis, we used either an average projection of one to five frames at the base of the crypt or at an upper layer about 15 µm closer to the villi. This upper layer was at approximate at the +3 cell position in the crypt^3^. Unless otherwise noted, displayed images are average projections of five frames and intensity adjustment was limited to linear scaling.

### Measurement of number of nuclei

In crypts labeled with Hoechst, we used projected images (2-5 frames in average) of crypt base processed as above. Each nucleus at the crypt base was numbered and individually tracked over time in each images session up to two hours. About 30 nuclei were visible and counted for each crypt.

### Rearrangement score for pattern disparity

To describe the alternating patterns of Paneth and Lgr5 cells, we devised a network graph representation using images of the crypt base. We manually defined nodes as the center of each Paneth cell. Two nodes were connected by edges if the two Paneth cells did not touch each other and a straight line between the two nodes did not touch any other Paneth cells. Edges represent the separation of Paneth cells by Lgr5 cells (Fig. 2c). The resulting network graph was used to calculate for a crypt with *n* Paneth cells an adjacency matrix at baseline, *M*_*baseline*_, and 1 day after ablation, *M_1_*. A rearrangement score that increases with more disparity between two patterns was defined as

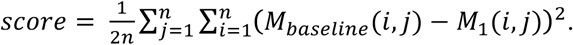

### Measurement of fluorescence intensity

To assess stability of Lgr5-GFP signals, we used processed and projected image (2-5 frames in average). Mean fluorescence intensity was measured from manually selected areas from same Lgr5-ISC and Paneth cell near the ablation at each time point. The analysis regions were selected by a user blinded to the treatment (control versus DBZ) and the timing of each image to reduce possible bias.

### PI label quantification and tracking

In animals injected with PI, projected images (2-5 frames average projection) from the processed stack were used from the crypt based and the upper layer. The area of PI-labeling was measured in ImageJ by manually drawing an outline of PI signal. To the track the position of the PI-labeled ablation debris along the crypt-villus axis as it moved into the crypt lumen, each image in the stack of images was processed with median filter to reduce noise. In order to maintain consistency in spacing, we did not delete moving frames. The first frame of the crypt base was used as the reference to align the different time points. The position of the lowest edge of the PI signal was recorded at each time. Linear regression was performed and plotted for each group.

### Tracking of Paneth cell movement within crypt base

To measure the movement of Paneth cells, we used images of the crypt base processed as above with both projections and single frames. Each Paneth cell at the crypt base was numbered and individually tracked. Using image J, the X-Y coordinates of the center of the Paneth cells were manually measured at baseline, 10, 30, 60 minutes after ablation. To align images between time points, we matched the position of center of the crypt so that the trajectory of each Paneth cell was identified (Suppl. Fig. 2e). The sum of the total distance moved by each Paneth cell was plotted using Graphpad software. The untreated group includes animals that were injected with saline as a sham treatment and animals with no injection, because saline injected, young mice resulted in similar dynamics as young, un-injected mice.

### Measurement of crypt lumen and outer circumference

The inner and outer circumference of the crypt was measured in single processed image frames. The circumference was measured by manual tracing along the lumen edge and the outer edge of Lgr5 labeling in ImageJ. Inner and outer circumference was measured from same image frame. Since crypts in animals with no injection did not show significant difference from sham, saline-injected animals, both data were included and displayed as the untreated group.

### Statistics

Graphpad Prism was used to prepared all graphs and for statistical analysis. Unless otherwise stated, to test for differences at various points in time, we used 2-way ANOVA with Tukey’s multiple comparison test and other tests are noted in text and captions.

## Acknowledgements

This study was supported by the Empire State Stem Cell Fund through New York State Department of Health Contract # C30293GG and NIH R01 GM114254. DMH is supported by NIA R56AG052981, P30AG038072 and the American Federation for Aging Research (AFAR).

## Author Contributions

J.C. developed methods, conducted experiments, wrote the manuscript and performed analysis. N.R., P.G., D.J.J. performed experiments and developed methods. S.M.L., T.T., D.M.H., X.S. and N.N. edited the manuscript and designed the experiments. N.N. supervised, designed experiments and wrote the manuscript.

## Additional Information

Competing Interests: The authors declare that there are no competing interests.

